# The Stochastic Pacemaker: Cumulative Behavioral Noise Drives Morphological Plasticity in Pea Aphids

**DOI:** 10.64898/2026.03.09.709630

**Authors:** Xiaomi Liu, Kristopher Murdza, Yuxu Feng, Leila Lin, Elizabeth I. Croyle, Jennifer A. Brisson

## Abstract

Phenotypic variation within a single genotype under the same environment (intragenetic variation), the biologically meaningful part of V_error_, is frequently treated as a statistical nuisance rather than a biological reality, yet it represents an evolutionary driver of fitness that remains poorly integrated into evolutionary theory. The mechanism that translates such stochasticity into deterministic developmental phenotypic outcomes is not well understood. Here, we test a cumulative stochasticity model using wing polyphenism of the pea aphid (*Acyrthosiphon pisum*), where asexual mothers produce winged instead of wingless offspring in response to tactile cues. The model predicts that stochastic variation in maternal locomotor behavior alters the rate of tactile cue accumulation and thereby influences the probability of producing winged offspring. We demonstrate that maternal locomotor activity acts as a "stochastic pacemaker", where an individual’s movement determines the rate at which it actively constructs its environment and accumulates environmental cues. Our results reveal that genotypes differ significantly in both mobility and the temporal pattern of wing induction, with behavioral variation explaining approximately 20% of the total phenotypic variance across genotypes. Crucially, we show that maternal mobility increases progressively during crowding, accompanied by significant temporal heteroscedasticity. This expanding variance and increase in mean suggest that initial, trivial stochasticity is magnified into systematic behavioral divergence through the integration of environmental signals. By demonstrating that total accumulated locomotor activity predicts offspring phenotype, we provide a mechanistic bridge between transient behavioral noise and stable morphological shifts. More broadly, our work reveals that V_error_ is a dynamic product of behavioral history, suggesting a fundamental role for individual-level niche construction in generating macro-phenotypic diversity.

## Introduction

Understanding the source of phenotypic variation has long been a key focus in evolutionary biology because this variation is the target of natural selection. From a quantitative genetic view, phenotype variance can be partitioned as V_P_ = V_G_ + V_E_ + V_GxE_ + V_error_, where each component represents a distinct source of variation (Lynch and Walsh, 1998; Tonsor et al., 2013; Westneat et al., 2015). Genetic variance (V_G_) captures differences among genotypes within a single environment, whereas environmental variance (V_E_) reflects phenotypic plasticity, the differences generated when genetically identical individuals experience different environments (Whitman and Agrawal, 2009). Genotype-by-environment interactions (V_G×E_) represent heritable differences in plasticity itself, providing another target for evolutionary change (Via and Lande, 1985; West-Eberhard, 2003a; Whitman and Agrawal, 2009). The final component, the residual term, V_error_, captures phenotypic differences among genetically identical individuals raised in nominally identical environments (Scheiner, 1993; Tonsor et al., 2013; Viney and Reece, 2013). Despite each component’s potential to impact evolutionary dynamics, they differ markedly in how well their underlying mechanisms are understood.

Of particular interest here is the understudied role of V_error_. This variation arises from several sources, such as measurement error, unmeasured or unmeasurable microenvironmental heterogeneity, and true stochastic developmental processes, like bet-hedging and stochastic gene expression (Kærn et al., 2005; Raj and Oudenaarden, 2008; Slatkin, 1974; Starrfelt and Kokko, 2012; Tonsor et al., 2013). The biologically meaningful portion of this residual variance, here referred to as intra-genotypic variation, can have a genetic basis and fitness consequences (Cai et al., 2008; Feinberg and Irizarry, 2010; Fraser et al., 2004; Kærn et al., 2005; Viney and Reece, 2013; Westneat et al., 2015). Yet despite its potential importance for evolution, intra-genotypic variation is frequently treated as a statistical nuisance rather than a biological reality, and has yet to be properly integrated in evolutionary theory (Cleasby and Nakagawa, 2011; Westneat et al., 2015). Characterizing this ‘residual’ component is essential for capturing the full spectrum of variation available to selection and for shedding light on understudied, interesting biological processes that shape phenotypic expression (Viney and Reece, 2013; Westneat et al., 2015).

Behavior provides a compelling candidate mechanism for generating such variation. As the most flexible response to environmental change, behavior serves as a rapid and dynamic interface between individuals and their environments (Duckworth, 2009; West-Eberhard, 2003b). Behavioral traits are highly variable, even among clonal individuals (Bierbach et al., 2017; Schuett et al., 2011), hinting that behavioral stochasticity may be a primary driver of residual phenotypic variance (V_error_). Furthermore, through niche construction, organisms, especially through behavior, can modify their own micro-environments, thereby altering the selective pressures and expression of non-behavioral traits (Duckworth, 2009; Odling-Smee et al., 2003, 1996). Despite long-standing theoretical predictions that behavior can modify the environment and thereby influence the expression and evolution of other traits (Bateson, 2004; Duckworth, 2009; West-Eberhard, 2003b), empirical studies that quantitatively link stochastic behavioral noise to non-behavioral variance remain comparatively limited. Identifying such connections would provide a critical, and currently missing perspective on how behavior contributes to the generation and evolution of phenotypic diversity.

Aphids provide an exceptional opportunity to empirically study the connection between behavioral variation to non-behavioral phenotypic outcomes. Their cyclical parthenogenesis enables the production of genetically identical individuals, allowing intra-genotypic variation to be isolated. Aphids also exhibit a well-characterized wing polyphenism, in which asexual females give birth to clonal daughters that are either winged or wingless depending on environmental conditions (Braendle et al., 2006; Brisson, 2010; Deem et al., 2024; Sutherland, 1969). Importantly, tactile stimulation generated during crowding is the key cue that induces more winged relative to wingless offspring production (Johnson, 1965; Müller et al., 2001; Sutherland, 1969; Sutherland and Mittler, 1971). Because tactile stimulation arises from physical interactions among individuals, variation in movement directly modulates cue exposure. We propose that individual locomotor behavior acts as an active, sensory-driven pace maker. Through their own movement, individuals, even clonal individuals, actively modulate the frequency of tactile stimuli they encounter, essentially "constructing" the environmental signal they receive. Thus, behavior is not only a consequence of an organism’s state, but also the causal driver of the developmental trajectory.

In this study, we used the pea aphid wing polyphenism to examine how both inter- and intra-genotypic maternal locomotor behavioral variation shape offspring phenotype compositions. We test the hypothesis that intra-genotypic variation (V_error_) is not an untraceable statistical constant, but a cumulative product of behavioral history. We propose a cumulative stochasticity model, where behavior acts as a "stochastic pacemaker": if locomotor activity facilitates tactile stimulation, then even minor initial differences in movement should undergo a process of linear accumulation over time. Our data demonstrate that genotypes differ significantly in both mobility and wing induction dynamics, with their correlation explaining about 20% of the total phenotypic variance. Crucially, within specific genotypes, individual mobility predicts winged offspring proportions, identifying behavior as a significant biological driver of V_error_. By quantifying the temporal increase in locomotor and its variance over time, we show that mobility increases progressively during crowding, magnifying initial stochasticity into systematic behavioral divergence. This provides a mechanistic bridge between stochastic behavioral noise and deterministic morphological outcomes. More broadly, our work addresses a critical gap in evolutionary theory by demonstrating how both inter- and intra-genotypic variation arises through behavioral integration and scales to generate macro-phenotypic diversity.

## Materials and Methods

### Aphid Stocks

Pea aphid (*Acyrthosiphon pisum*) stocks were reared on *Vicia faba* seedlings (Improved Long Pod, Harris Seeds) by transferring four first instars on fresh seedlings every 11-13 days. Each plant was grown in a single plastic plant pot (S17648, Fisher Scientific) and enclosed in a cylindrical cage cut from clear PETG tubes (VISIPAK) sealed with fine mesh netting (Noseeum netting, 117 inches, Barre Army Navy Store). Cages were maintained in an incubator under long-day conditions (16L:8D), with a relative humidity of 30 ± 10% and a temperature of 19 ± 3.5°C. These environmental conditions enable the pea aphid to remain in an asexual life stage and reproduce viviparously. Stocks were reared under the described conditions for multiple generations to eliminate transgenerational density effects before the experiment.

### Recording Setup

Four aphid genotypes were used: ROC1, SSC3, 375, and 663. Lines ROC1 and 375 show a pink color morph, while lines SSC3 and 663 show a green color morph (Figure 1A). To minimize stimuli from outside the recording, fresh adult aphids were collected directly from plants and immediately used for behavioral recording. Due to the limited ability of existing tracking software to reliably differentiate between visually similar moving individuals, particularly when animals frequently interact or obstruct one another, individual identity error is unavoidable (Dell et al., 2014). Here, we leveraged the natural color variation of pea aphids to ensure 100% identification accuracy. During the recording, one focal aphid of one color morph, either pink or green, was paired with 11 background aphids of the alternate color. Together, 12 aphids were placed in a 35 mm × 10 mm Petri dish. Six such Petri dishes were simultaneously recorded on a 5000K 70% brightness video light (RALENO, PLV-S116) for 7hrs using a mobile phone camera (Figure 1B). For the recording, line ROC1 was paired with line SSC3 and is hereafter referred to as pair 1 pink and pair 1 green, respectively. Similarly, lines 375 and 663 were paired and are referred to as pair 2 pink and pair 2 green. Following the recording, the focal aphids were individually transferred onto fresh seedlings and allowed to reproduce for 24 hours. The winged offspring percentage was scored upon adulthood. This experimental design enabled the accurate collection of behavioral and phenotypic data for each focal aphid, eliminating the risk of mismatches between behaviors and phenotypes due to limitations in tracking accuracy. We also collected an additional 3-hour recording for ROC1 (pair1 pink) for the reasons listed in the later context. All recordings started around 9 a.m..

**Figure 1.**
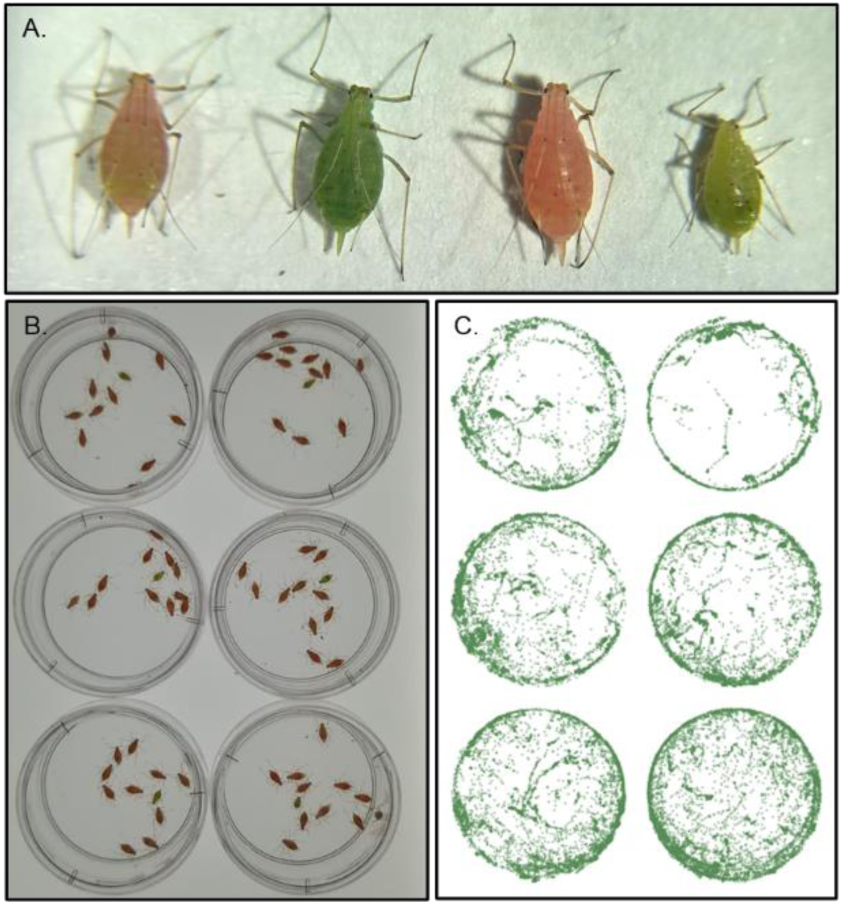
Experimental setup and visualization of behavioral recordings. (A) Representative photo of the four pea aphid genotypes used in the study. From left to right: pair 1 pink, pair 1 green, pair 2 pink, and pair 2 green. (B) Representative single frame from a behavioral recording showing aphids within the observation arena. (C) Overlay of all detected aphid centroid positions across the full recording period for one example recording, with 1 position sample every 2s, illustrating spatial coverage of aphid movement within the arena.

### Video Analysis

To estimate the amount of tactile stimulation aphids received during the recording period, we directly counted interactions. Two observers independently quantified interaction to account for human error. Due to time and resource limitations, we collected only interaction data for the pair 1pink recording.

To estimate aphid activity levels, we extracted each aphid’s movement data for downstream analysis. One frame was extracted every two seconds from the original recording with FFmpeg (Tomar, 2006). Frames were then cleaned and processed using customized scripts and were subsequently split into pink and green channels using the ColorDeconvolution2 plugin (Ruifrok and Johnston, 2001) in ImageJ (Schneider et al., 2012). Frames from the color channel containing focal aphids were further tidied up with customized scripts and used to track aphid movement using TrackMate (Ershov et al., 2021; Tinevez et al., 2017). The tracking results were exported as data tables containing X and Y coordinates for aphid centroid positions in each frame. All positions for a recording were plotted to determine the boundary for each Petri dish (Figure 1C). Boundaries were assigned to different Petri dishes and corrected for false detection using customized scripts. After false detection correction, each Petri dish was corrected for camera distortion by resetting x boundaries to range from 0 to 400 (pixels) for both X and Y coordinates. Frames from the color channel that contain background aphids were tracked with StarDist (Schmidt et al., 2018). The trajectory results were exported as data tables with trajectory distance for each identified track.

### Profiling temporal dynamics in response to crowding

To test the effect of different crowding durations on offspring wing percentage, we crowded aphids for varying durations: 0 hours (no crowding), 4, 7, 10, or 24 hours. All treatments started around 8 a.m. During the crowding, 12 healthy adult aphids were placed in a 35 mm Petri dish and then stored in an additional 90 mm Petri dish. A sheet of damped Waltman paper was placed inside both the 35 mm dish and the larger Petri dish to prevent the aphids from severe dehydration. After crowding, aphids were transferred individually onto fresh seedlings and allowed to reproduce for 24 hours. For the 0hr group, aphids were collected directly from the plants and placed on new seedlings for 24hrs to reproduce. The offspring wing percentage was scored upon adulthood. Following the initial data collection, based on the results, we selectively collected additional data at 1, 2, and 3hrs for pair 1 pink and at 36 hours for pair 2 green.

### Data analysis

We only included records in which treated aphids produced more than two offspring. For both focal and background aphids, we measured activity level by calculating the total distance the aphid moved. For the focal aphids, the distance was calculated by summing up the geometric distance traveled between every frame. For the background aphids, the total travel distance was calculated by summing up the distance for all detected trajectories. Activity data, interaction number, and offspring wing percentage data were all tested for normality with the Shapiro test. Because some data were not normally distributed (Table S1), we used nonparametric tests or rank-transformed data when needed. When multiple comparisons were performed, we adjusted p-values with the Benjamini-Hochberg method (Benjamini and Hochberg, 1995).

All analyses were done in R 4.3.1.

## Results

### Locomotor activity strongly predicts tactile stimulation rate

To lay the foundation for downstream hypotheses and analyses, we first tested the expectation that greater mobility would increase the likelihood of physical interaction between aphids, thereby enhancing tactile stimulation. We measured interaction number as a direct indicator of tactile stimulation, and quantified activity level using total travel distance as a proxy for mobility. As expected, we observed a tight correlation between the number of interactions and travel distance (Figure S1, Table S1, Spearman Rank Correlation Test, Observer 1: correlation coefficient ρ = 0.883, adjusted p-value = 1.101 × 10^-12^; Observer 2: correlation coefficient ρ = 0.893, adjusted p-value = 4.948 × 10^-13^). This result suggests that more active aphids experience greater tactile stimulation, providing a basis for subsequent analyses examining the relationship between activity and plasticity response.

### Intra-genotypic variation: individual behavioral differences impact phenotypic variation

To investigate the source of V_error_, we hypothesized that individual-level activity, which varies even among clonal individuals, introduces microenvironmental heterogeneity that contributes to final phenotype variation. We measured the final offspring wing percentage phenotype and individual locomotion, and performed correlation tests within each genotype. Significant correlations between offspring wing percentage and activity level were detected in two out of the four tested genotypes (Figure 2, Kendall Rank Correlation Test, pair 1 green adjusted p-value = 0.024, pair 2 pink adjusted p-value = 0.024). We then estimated how much of the phenotypic variance could be explained by behavior. We calculated Kendall’s tau^2^ as a nonparametric proxy of association. For pair 1 (pink) and pair 2 (green) where significant correlations were detected, the tau² values were 0.115 and 0.097, respectively. We also applied a linear regression model, recognizing that the data did not follow a normal distribution; thus, interpretation should be cautious. We obtained R^2^ values of 0.248 and 0.177 for pair 1 pink and pair 2 green, respectively. The higher R² and lower tau^2^ suggest a generally linear yet noisy relationship with tied ranks, consistent with the nature of our data (Figure 2). Taken together, these results indicate that intra-genotypic individual-level activity accounts for a considerable yet incomplete portion of final phenotype variation, helping to identify part of the source of the V_error_ term.

**Figure 2.**
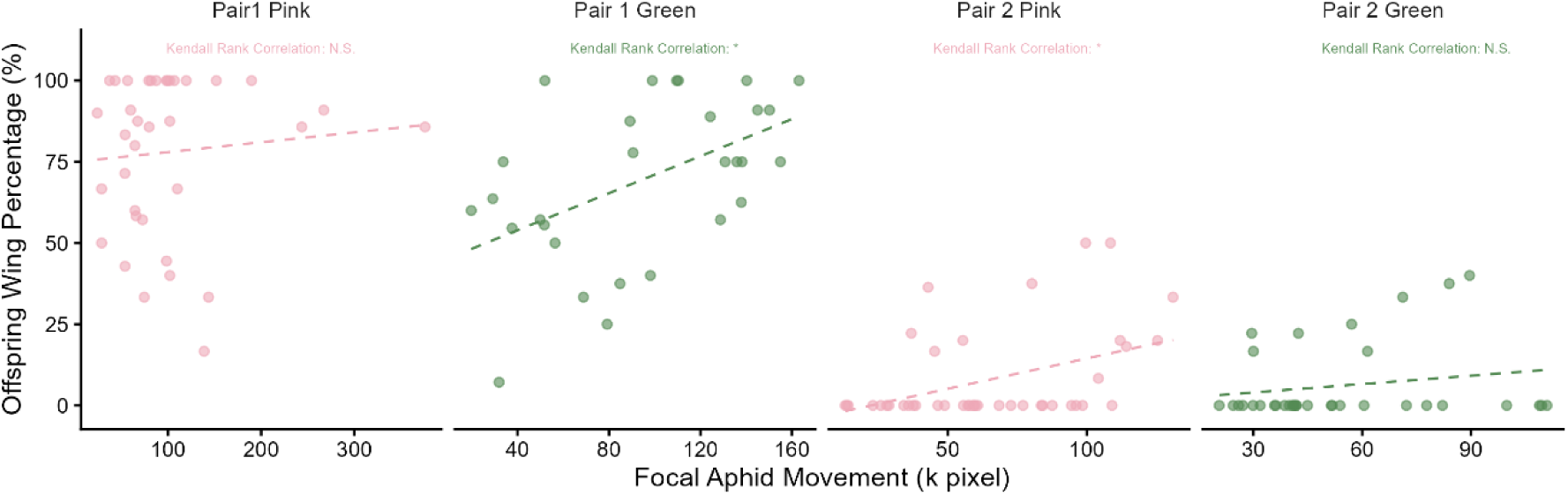
Intra-genotypic variation in maternal locomotor behavior correlates with offspring wing phenotype in some genotypes. Scatter plots show the relationship between focal aphid locomotor activity (x-axis) and the percentage of winged offspring (y-axis). Each panel shows a single pea aphid genotype, with each point representing one focal individual.

### Inter-genotypic variation: behavioral divergence across genotypes contributes to phenotypic variation

To assess inter-genotypic variation in the wing percentage phenotype, we collected offspring wing percentage data post-crowding and conducted a nonparametric statistical test. We found that pair 1 pink and green have a higher propensity to produce winged offspring compared to pair 2 pink and green (Figure 3A, Table S1, Kruskal-Wallis chi-squared = 105.9, p-value = 8.3 x 10^-23^, see detailed Dunn test result in Table S1), supporting the idea that there are genetic differences underlying this wing induction phenotype. The four tested genotypes also showed non-homogeneous levels of data dispersion (Figure 3A, Table S1, Fligner-Killeen test, p-value = 2.797 × 10^-6^), suggesting that the variation level for the phenotype itself also has a genetic basis.

**Figure 3.**
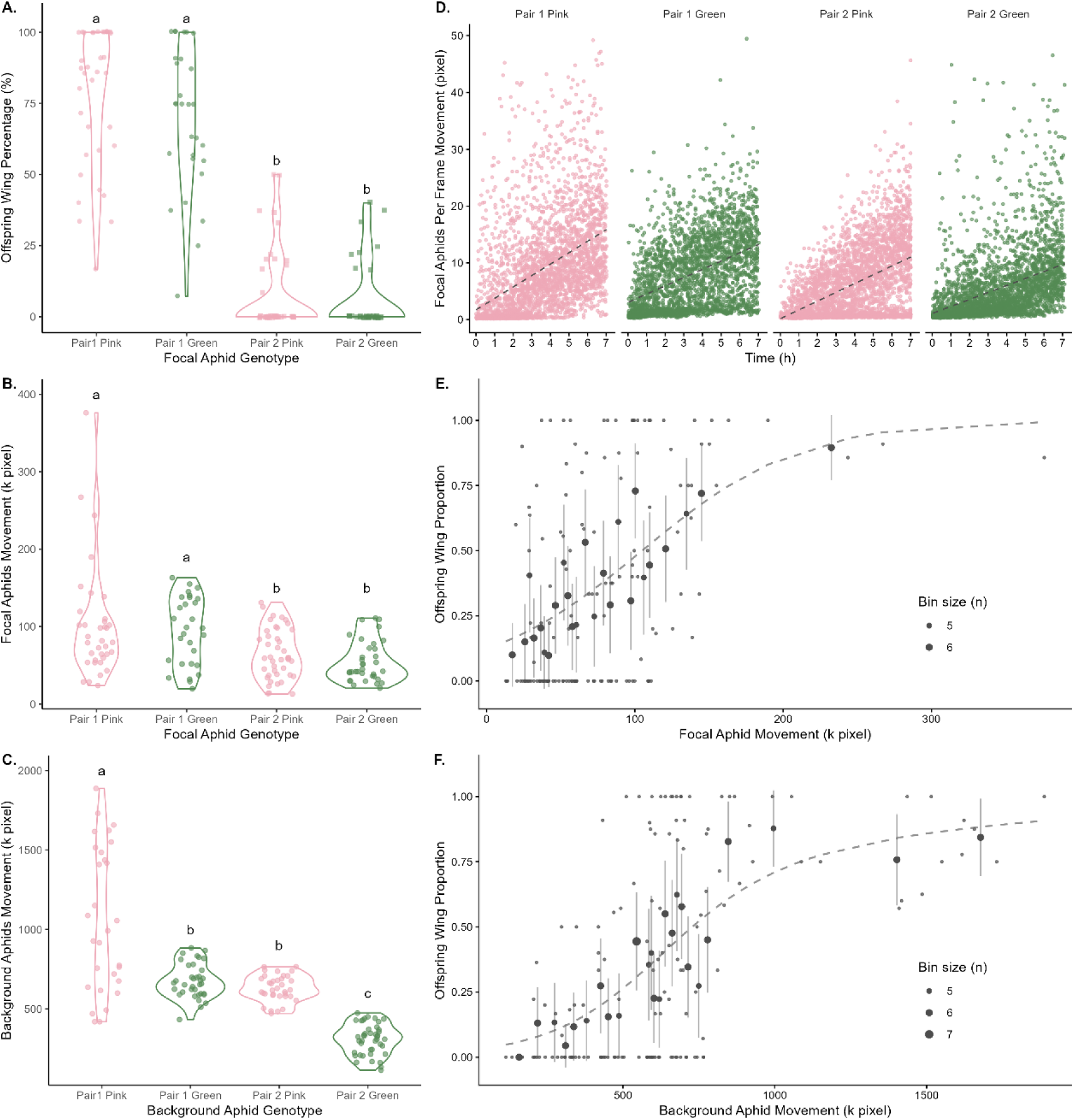
Inter-genotypic variation in maternal locomotor behavior contributes to the offspring wing phenotype. (A) Offspring winged percentage produced by focal aphids from each of four genotypes. (B) Locomotor activity of *focal* aphids from each of four genotypes. (C) Locomotor activity of aphids from the same genotypes when serving as *background* individuals. D) Temporal profile of locomotor activity, showing per-frame movement averaged across 5 min time windows (y-axis) as a function of time during recording (x-axis). Colors represent aphid color morphs; each panel represents one of the four genotypes. (E) Association between *focal* aphid locomotor activity and *focal* aphid offspring wing proportion. (F) Association between *background* aphid locomotor activity and *focal* aphid offspring wing proportion. For panels E and F, the raw data points are shown as small grey points. Data were grouped into 25 bins along the x-axis (distance moved), with uneven bin widths to yield approximately equal sample sizes per bin (5–7 points). Black points represent the mean value for each bin, with the light gray lines indicating the binomial standard error. GAM model fits are shown as light grey dashed lines.

To assess intergenotypic variation in behavior, we calculated the total geometric distance traveled by aphids, both as focal and background individuals, and conducted nonparametric statistical tests. When acting as the single focal aphid, both pair 1 pink and pair 1 green showed significantly higher activity levels compared to the two genotypes from pair 2 (Figure 3B, Kruskal-Wallis chi-squared = 20.800, p-value = 1.158 x 10^-4^, see detailed Dunn test results in Table S1). When comparing the activity level of the background aphids, we found some similarity in behavior patterns, where pair 1 pink moved significantly more than all other genotypes (Figure 3C, Kruskal-Wallis chi-squared = 95.107, p-value < 1.751 x 10^-20^, see detailed Dunn test results in Table S1). However, pair 1 green now showed a similar activity level to pair 2 pink (Figure 3C; see detailed Dunn test results in Table S1). Pair 2 green showed the lowest activity level (Figure 3C). The movement of focal and background aphids cannot be directly compared because they were collected using different tracking methods. Additionally, background aphid travel distance represents the summed activity of all eleven individuals within a Petri dish, reflecting summary statistics to some extent and resulting in a narrower data dispersion. For both focal and background travel distance, pair 1 pink exhibited greater variation than the other three genotypes (Fig. 3B, 3C, Fligner-Killeen test, focal aphids distance p-value = 0.015, background aphids distance p-value = 1.093 × 10^-13^), suggesting increased individual-level behavioral variation in this genotype. Together, our results show rich inter-genotypic variation in wing phenotype and behavior.

We next examined how locomotion changed over time during the recording period. Mean travel distance was calculated for each dish (1-6) and recording date using 5-min windows. Time was standardized (mean-centered and scaled by its standard deviation) to match the magnitude of speed and to improve numeric stability, yielding approximately 2500 – 3500 observations per genotype. Mean travel distance was analyzed using a linear mixed-effect model, where time, genotype, and interaction terms were included as fixed effects. Individual identity was modeled with both random intercept and slope to account for repeated measurements and individual trajectory. Residual variance was allowed to vary as a function of time to test for heteroscedasticity. As shown in Figure 3D, time had a strong positive impact on locomotion. Genotype-specific slopes were estimated as follows: pair 1 pink (β = 4.10, 95% CI 3.23 - 4.98), pair 1 green (β = 2.84, 95% CI 1.88 - 3.80), pair 2 pink (β = 3.32, 95% CI 2.51 - 4.13), and pair 2 pink (β = 2.69, 95% CI 1.81 - 3.56), with all estimated slopes significantly greater than 0. This result indicates a progressive increase in locomotion throughout the assay. The time × genotype interaction was significant for some genotypes (Table S1), demonstrating genotype-dependent differences in the magnitude of temporal response relative to the reference genotype (pair 1 pink). Thus, genotypes differed not only in baseline mobility but also in the rate of behavioral change over time. Including individual-specific random slopes significantly improved model fit compared to a random-intercept-only model (likelihood ratio test, p < 0.0001; Table S1), indicating substantial among-individual variation in temporal trajectories. Furthermore, allowing residual variance to scale with time significantly improved model fit relative to a homoscedastic model (likelihood ratio test, p < 0.0001; Table S1, Figure 3D), demonstrating an increase in within-genotype locomotor variability over time. Taken together, these results revealed a strong temporal increase in locomotion behavior, as well as inter- and intra-genotypic variation in movement dynamics.

After observing substantial variation within and among genotypes, we next examined whether the relationship between activity and phenotype holds when all genotypes are considered simultaneously. To do this, we modeled offspring wing percentage and activity data from the four genotypes combined. Given that our data were enriched for 0% and 100%, we converted the percentages to proportions and employed a generalized additive model with a quasibinomial family. The analysis revealed a significant positive linear relationship between offspring wing percentage and focal aphid movement (Figure 3E, p-value = 9.685 × 10^-7^, effective degrees of freedom = 1.000). A similar significant but non-linear relationship was detected with the background aphid movement data (Figure 3F, p-value = 0, effective degree of freedom = 2.016). Additionally, the focal aphid movement explains 17.1% deviance, while the background aphid movement explains 29.5% deviance (Table S1). Taken together, our results suggest a general pattern: more active individuals produce a higher proportion of winged offspring, implying that inter-genotypic behavioral variance can account for approximately 20% of inter-genotypic phenotypic variance.

### Genotype-specific Behavior-Sensitive Window

As we observed a general significant correlation between behavior and phenotype across genotypes, and in two out of four genotypes, we next asked why we failed to detect such a correlation in pair 1 pink, which nonetheless showed substantial variation in both behavior and phenotype. We hypothesized that different genotypes exhibit distinct dynamics in generating the plasticity response, thus affecting the behavior-sensitive window. To test this, we measured wing percentage data across varying time (0, 4, 7,10, or 24hrs) spent crowded to capture temporal dynamics. Our results revealed drastically different responses among genotypes (Figure 4A). Pair 1 pink responded rapidly to crowding, reaching a plateau after only 4 h, whereas pair 1 green and pair 3 pink reached their maximal responses after approximately 7 h and 10 h, respectively. In contrast, pair 2 green exhibited a general lack of response. To refine our temporal resolution, we collected additional early time points (1 h, 2 h, and 3 h) for pair 1 pink and an extended 36 h time point for pair 2 green to maximize signal detection, while maintaining the response shape. The maximum response at the plateau also differed among genotypes, with pair 1 pink and pair 1 green showing a high propensity for producing winged offspring, whereas pair 2 pink showed a moderate to low propensity (Figure 4A). Overall, these results demonstrate that time is a strong positive effector of the final phenotype in three out of four tested genotypes, supporting our hypothesis (Figure 4A, Kruskal-Wallis Test, adjusted p-value pair 1 pink: 5.544 × 10^-11^, pair 1 green: 1.401 × 10^-4^, pair 2 pink: 6.932 x 10^-5^). The early saturation of the response in pair 1 pink explains the absence of a detectable behavior-phenotype correlation under the 7 h recording setting, as the phenotype was already at its plateau. Contrastingly, the lack of a correlation in pair 2 green likely comes from its limited overall responsiveness to crowding. As for pair 1 green and pair 2 pink, the 7 h window corresponds to the dynamic range in which variation in activity generates variation in tactile stimulation, resembling either shorter or longer crowding times, thus affecting the final phenotype output. Taken together, these results indicate that genotypes differ in their plasticity response dynamics, revealing inherent sensitive windows in which behavior-phenotype correlations can be detected.

**Figure 4.**
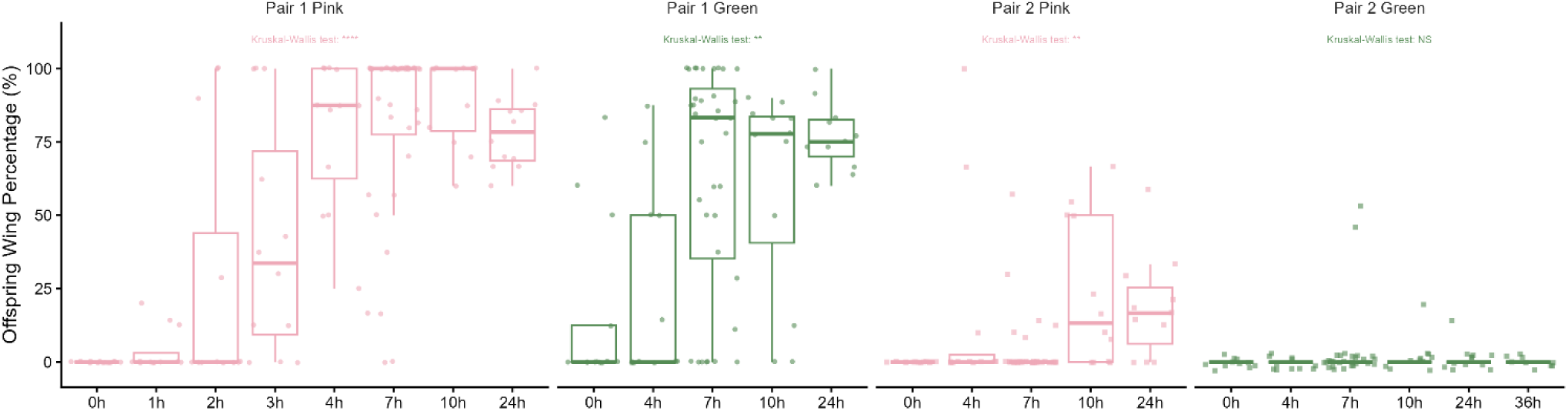
Genotypes exhibit distinct temporal patterns of wing induction in response to crowding. Relationship between crowding duration (x-axis) and offspring wing percentage (y-axis). Each small panel corresponds to each of the four genotypes.

Based on the unique behavior-sensitive window hypothesis, we predicted that a significant correlation between phenotype and behavior would be detected at an earlier time window for pair 1 pink. To test this, we repeated the recording at 3 h for pair 1 pink. The analysis detected a significant correlation between movement and offspring wing percentage, supporting our hypothesis (Figure 4B, Kendall Rank Correlation Test, p-value = 0.016). Similar to pair 1 green and pair 2 pink, 7h recording data, the tau2 is 0.093, suggesting a similar amount of variation is explained by intra-genotypic individual-level behavioral differences.

## Discussion

A central contribution of this study is the empirical demonstration that maternal behavior serves as a "stochastic pacemaker", explaining a substantial portion of offspring wing plasticity both within and between pea aphid genotypes. At the intra-genotypic level, we show that seemingly stochastic differences in maternal movement create heterogeneous microenvironments by modulating active niche construction, altering the rate at which individuals encounter and accumulate tactile cues, which in turn alter the likelihood of producing winged offspring. In this framework, variance previously classified as stochastic residual error (V_error_) is not random noise, but rather an integrated consequence of each mother’s behavioral history. Behavioral stochasticity, therefore, becomes biologically consequential through cumulative signal acquisition. At the inter-genotypic level, the strong correlation between maternal behavior and offspring phenotype across genotypes indicates that heritable differences shape how mothers perceive, integrate, and respond to environmental cues. Importantly, genotypes differed not only in mean mobility and wing plasticity but also in the temporal dynamics of behavioral change under crowding. These differences suggest that selection can act not only on the magnitude of wing plasticity but also in the temporal dynamics of signal integration. Together, linking transient behavioral variability to stable morphological outcomes, our results provide empirical support for the cumulative stochasticity model and position behavioral variance as an evolvable component of developmental plasticity.

Under the Cumulative Stochasticity Model, maternal behavior modulates morph determination through iterative amplification of tactile cues. Tactile stimulation generated by movement promotes additional movement, gradually magnifying small initial differences in activity into substantial divergence in cue exposure over time. Behavioral responses can also potentially integrate other environmental inputs. For example, alarm pheromone ((E)-β-farnesene) induces "pseudo-crowding" by rapidly increasing walking behavior, which in turn elevates the proportion of winged offspring (Kunert et al., 2005). Similarly, intensified foraging behavior is observed under starvation, another nutrition-associated condition that promotes offspring wing production (Baz et al., 2025). Such behaviorally mediated cue amplification is widespread across insects. Because behavior is typically the earliest response to environmental change and is itself one of the most plastic trait (Dingemanse and Wolf, 2013; Duckworth, 2009; Gosling, 2001; Sih et al., 2004; Wolf et al., 2008; Wolf and Weissing, 2012), it can act as a primary modifier that strengthens or prolongs initial cues before any downstream physiological development pathway are involved. Locust phase polyphenism provides a classic example of behavior reinforcement: solitary individuals increase aggregation behavior within hours of crowding, and this behavior then reinforces the gregarizing stimuli (Bouaïchi et al., 1995; Ellis, 1963; Gillett, 1973; Roessingh and Simpson, 1994; Rogers et al., 2003). Consistent with this general pattern, our data (Figure 3D) shows progressively increasing maternal activity as crowding proceeds. This temporal escalation supports the interpretation that behavior does not merely respond to crowding but actively intensifies cumulative signal exposure, thereby strengthening its contribution to morph determination.

Our findings integrate a model linking behavior, signaling dynamics, and offspring phenotypic plasticity. In this framework, mothers perceive tactile stimulation during crowding and transduce this information into signaling pathways, including glutamate, dopamine, ecdysone, and insulin/insulin-like growth factor signaling, that regulate wing plasticity (Brisson et al., 2026; Deem et al., 2024; Grantham et al., 2020; Liu and Brisson, 2023; Vellichirammal et al., 2017; Yuan et al., 2025, 2023). Behavioral variation potentially influences this system in two ways. First, differences in accumulated movement alter the intensity and frequency of tactile stimulation (Figure S1), thereby likely generating variation in signaling-molecule concentrations within and across genotypes. To evaluate how broadly such behavioral effects may apply, we examined multiple genotypes to increase the generality of our intra- and intergenotypic conclusions. Consistent with our intra-genotypic conclusion, another study using a different aphid genotype also reported a positive association between intra-genotypic behavioral activity and winged offspring production in a 2-individual, 2-h crowding setup (Yuan et al., 2025). Second, variation in the timing of the behaviorally sensitive window may interact with genotype-specific signaling thresholds to determine developmental outcomes, a hallmark of polyphenic systems (Nijhout, 2003, 1999; Pfennig, 2021). The interplay between behavioral modulation of cue exposure, genetically determined thresholds, and the temporal dynamics of signal production likely all contribute to the observed variation in winged-offspring production.

Despite considerable residual variance (V_error_) being explained by inter- and intra-genotypic behavioral variation, a large proportion of the variance remains unexplained. One source of developmental stochasticity for V_error_ can be bet hedging, a strategy favored when environmental cues are unreliable predictors of future conditions (Slatkin, 1974; Starrfelt and Kokko, 2012; Tonsor et al., 2013). Pea aphids, like many other systems, employ a mixed plasticity and bet-hedging strategy: mothers stochastically produce both morphs regardless of environmental conditions, ensuring that some offspring match unpredictable environments and thereby increasing overall survival of the species (Grantham et al., 2016). Such intrinsic stochasticity can blur the quantitative relationship between cue intensity and phenotypic outcome.

Finally, our results provide empirical support for the theoretical expectation that behavioral variation can shape, and in some cases even lead, the evolution of other traits (Duckworth, 2009; Odling-Smee et al., 2003, 1996; West-Eberhard, 2003b). In our system, maternal locomotion does not merely respond to environmental conditions; it modulates the effective strength and duration of crowding cues experienced by the individual. In doing so, behavior alters the local developmental environment that determines offspring morphology. This aligns with niche construction theory, which emphasizes that organisms actively modify their environments and thereby alter the selective pressures they experience (Odling-Smee et al., 2003, 1996). Under this framework, variation in movement rate translates directly into variation in cumulative signal exposure, thereby influencing the probability of wing induction. As we show, even clonal individuals present phenotypic variation rooted in maternal behavior history, demonstrating that behavioral variation can generate structured developmental divergence without genetic differences. More broadly, our findings suggest that individual-level micro-environmental heterogeneity may represent an underappreciated mechanism through which behavior contributes to macro phenotypic diversity.

---

**Figure S1.**
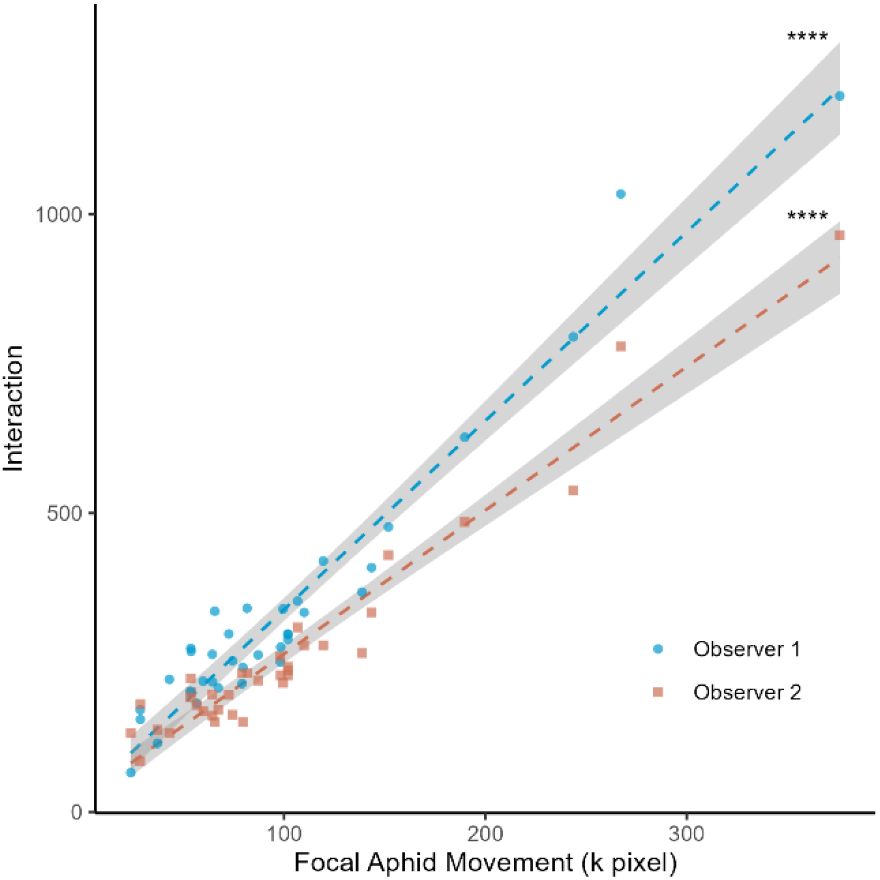
Tight correlation between aphid locomotor activity and interaction frequency. Interaction number (y-axis) is plotted as a function of focal aphids locomotor activity (x-axis). Colors and point shapes represent two independent observers who quantified the aphid-aphid interactions.

Table S1. Detailed statistical analysis table.

## Acknowledgements

This research is supported by the National Institute of General Medical Sciences within the National Institutes of Health under award number R35GM144001 to JAB. We thank Arron L. Forbes for helpful discussions and assistance in designing the interaction-counting criteria used in this study.

## Notes

### Competing Interest Statement

The authors have declared no competing interest.

### Summary of Updates

The title page has been updated to reflect the correct manuscript and author details. Author affiliation information has been corrected where necessary to ensure accurate institutional attribution, and the relevant ORCID identifier has been added and properly linked. In addition, the acknowledgements section has been updated to accurately recognize funding sources and contributions. No changes were made to the scientific content, analyses, or conclusions of the manuscript.

