## Supplementary figures and images for "The Stochastic Pacemaker: Cumulative Behavioral Noise Drives Morphological Plasticity in Pea Aphids"

### Figure S1

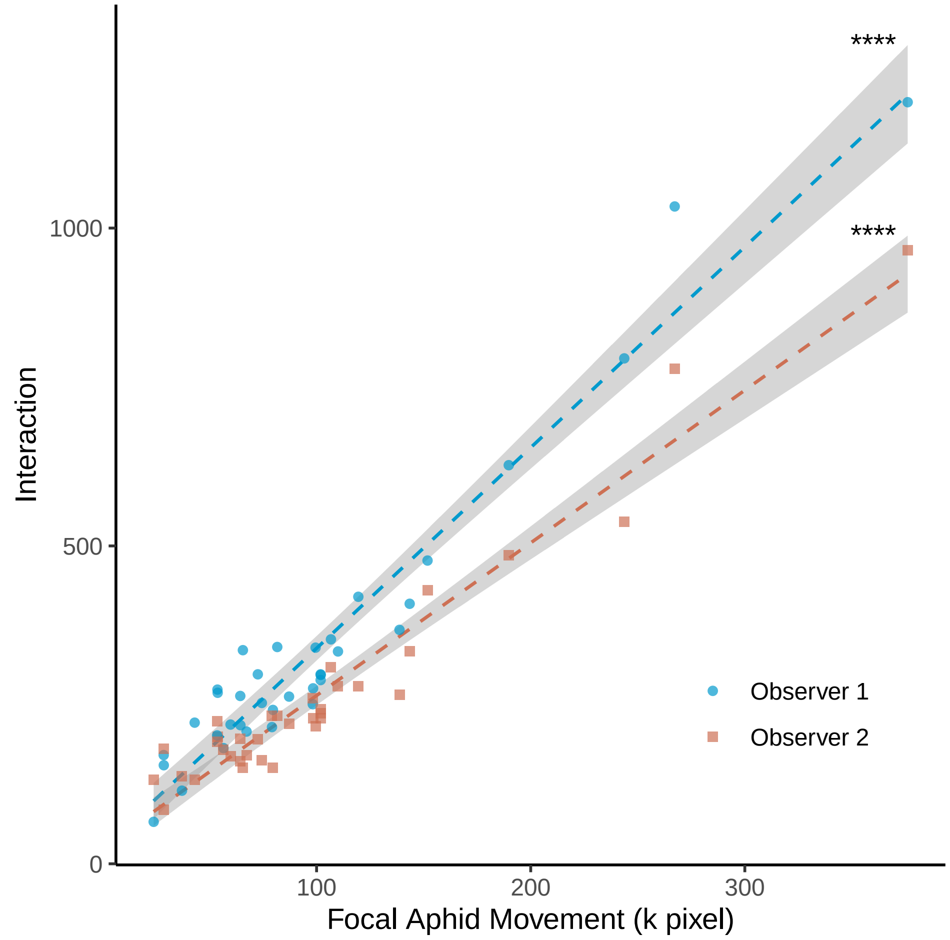
